# The evolution of color naming reflects pressure for efficiency: Evidence from the recent past

**DOI:** 10.1101/2021.11.03.467047

**Authors:** Noga Zaslavsky, Karee Garvin, Charles Kemp, Naftali Tishby, Terry Regier

## Abstract

It has been proposed that semantic systems evolve under pressure for efficiency. This hypothesis has so far been supported largely indirectly, by synchronic cross-language comparison, rather than directly by diachronic data. Here, we directly test this hypothesis in the domain of color naming, by analyzing recent diachronic data from Nafaanra, a language of Ghana and Côte d’Ivoire, and comparing it with quantitative predictions derived from the mathematical theory of efficient data compression. We show that color naming in Nafaanra has changed over the past four decades while remaining near-optimally efficient, and that this outcome would be unlikely under a random drift process that maintains structured color categories without pressure for efficiency. To our knowledge, this finding provides the first direct evidence that color naming evolves under pressure for efficiency, supporting the hypothesis that efficiency shapes the evolution of the lexicon.

## 1. Introduction

A substantial body of research suggests that languages are shaped by efficient communication (see e.g. Gibson et al., 2019 for a recent review). On this view, language evolution is driven, at least in part, by a functional need for communication to be both accurate and simple. This general idea has been pursued with respect to a number of specific aspects of language, including semantic categories (e.g., Kemp et al., 2018), with color naming as a prominent example. Many empirical findings suggest that languages tend to acquire new color terms with time, resulting in increasingly fine-grained color naming systems (Berlin and Kay, 1969; Kay and Maffi, 1999; MacLaury, 1997; Levinson, 2000; but see also Haynie and Bowern, 2016). More recently, it has been claimed (e.g., Lindsey et al., 2015; Regier et al., 2015; Gibson et al., 2017; Kemp et al., 2018; Zaslavsky et al., 2018; Conway et al., 2020) that this historical evolutionary process, and color naming more generally, are shaped by a need for efficient communication.

However, most research concerning the evolution of color naming has been based indirectly on synchronic cross-language comparison, rather than directly on fine-grained diachronic data collected in the field. There are some approaches that have approximated this ideal: e.g. Biggam (2012) considered historical texts; Kay (1975) considered informant age as a proxy for change over time; and Haynie and Bowern (2016) used phylogenetic methods to infer the history of color naming in a particular language family. However, these remain approximations: historical texts, while providing genuinely diachronic data, do not support analyses at a fine-grained level close to color perception; informant age is a reasonable proxy for change over time, but still a proxy; and phylogenetic reconstruction provides an inferred historical record rather than a directly measured one. In a recent exception to this general trend, Huis-man et al. (2021) explored the evolution of color naming in Japonic languages by directly comparing fine-grained data collected in the field at different points in time. Still, this approach remains unusual, and to our knowledge no prior study has used fine-grained diachronic data from the field with a view to examining questions of efficiency in the evolution of color naming.

Here, we do that. Specifically, we explore the role of efficiency in color naming evolution by considering fine-grained diachronic data from the field for a single language, Nafaanra (iso:nfr, Senufo, Ghana). We do this in a theory-driven manner, by testing quantitative predictions for language change previously derived from the theoretical framework of Zaslavsky, Kemp, Regier, and Tishby (2018, henceforth ZKRT). This framework integrates the proposal that languages evolve under pressure for efficient communication together with the Information Bottleneck principle (Tishby et al., 1999), which can be formally derived from rate-distortion theory (Shannon, 1959; Berger, 1971), the branch of information theory that characterizes optimal data compression under limited communicative resources.

We find that: (1) color naming in Nafaanra has changed during the recent past by adding new color terms and becoming more semantically fine-grained; (2) this has happened in a way that is consistent with pressure for efficiency as predicted by ZKRT; and (3) this outcome would be unlikely under a process of random drift that maintains structured color categories without pressure for efficiency. To our knowledge, this is the first finding that directly supports the proposal that color naming evolves under pressure for efficiency. Xu et al. (2016) previously used a related theoretical framework to show that a specific mechanism of semantic change — semantic chaining — shows signs of pressure for efficiency in a different semantic domain, that of names for containers. Our present work shows direct pressure for efficiency in language change that is not restricted to chaining, using a different framework that suggests a continuous evolutionary process (Zaslavsky et al., 2018; Zaslavsky, 2020), and in a domain — color naming — for which questions of evolution and language change have long been theoretically central.

In what follows, we first discuss color naming in Nafaanra, comparing data from 1978 with data that one of us (K.G.) collected in 2018. We then review the theoretical framework of ZKRT and test its predictions in the case of semantic evolution in Nafaanra color naming. We conclude by discussing implications of our findings.

## 2. Color naming and its evolution: The case of Nafaanra

Nafaanra is a Senufo language spoken in Ghana and Côte d’Ivoire, with approximately 61,000 speakers across all dialects (Simons and Gordon, 2006). The Nafaanra data in this study were collected in the town of Banda Ahenkro, Ghana. Community members estimate that the greater Banda region currently has around 20,000 speakers of Nafaanra spread throughout the area, with around 6,000 speakers in Banda Ahenkro proper (Garvin, 2017).

In Banda Ahenkro, Nafaanra is the most commonly spoken language and is used across all domains. However, within the Banda Ahenkro community, there are no known monolingual speakers of Nafaanra except for small children, as many Nafaanra speakers also speak Twi (iso:twi, Kwa, Ghana), a member of the Kwa language family (Simons and Gordon, 2006), and English, to varying degrees of frequency and fluency. Twi serves as a lingua franca beyond Banda Ahenkro, and English is the national language, learned and used in education. Proficiency for Twi is generally higher than for English, and Twi is used more frequently and across more domains. However, media is often in English, and thus, while proficiency in English is lower, exposure to English is still high. Despite the influence of Twi and English, Nafaanra is dominant for Nafaanra speakers in the Banda Ahenkro region. Community members understand the current overall language usage profiles to be comparable between 1978 and 2018 (the two years of data collection), with Nafaanra as the dominant language, and some Twi and English usage in trade and education respectively; however, speakers also report an increase in usage and exposure to Twi and especially English since 1978. One major factor in the increase in exposure is a change to both technology and lifestyle in the community. First, more community members now have access to television, which in particular has increased exposure to English. In addition, it is more common among younger generations for children to leave the community in the later years of schooling to receive education, and subsequently, to find a job, rather than pursuing agriculture, which was once the dominant occupation in the community. Outside of the community, people are exposed in particular to more Twi, and changes in education and occupational trends results in more exposure to English.

Color naming data for Nafaanra were initially collected in 1978 in Banda Ahenkro, as part of the World Color Survey (WCS; Kay et al. 2009), following WCS protocol.^1^ Participants in the WCS were shown each of the 330 color chips in the color naming grid shown in Figure 1, in a fixed random order, and asked to provide a name for each color. A total of 29 Nafaanra speakers participated in the 1978 survey, and the resulting data are shown in Figure 2A. The Nafaanra color naming system of 1978 is a 3-term system, with terms for light (‘fiηge’), dark (‘wͻͻ’), and warm or red-like (‘nyiε’).

**Figure 1.**
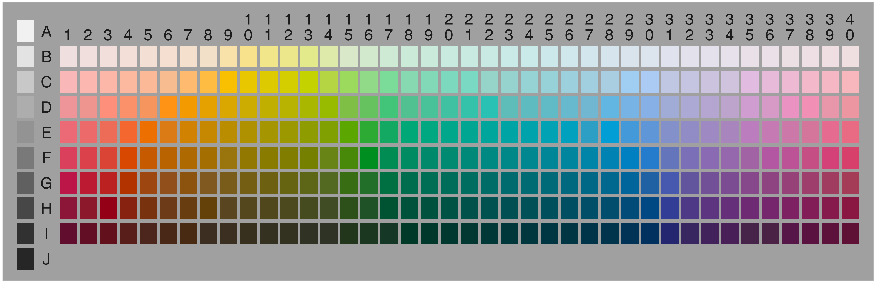
Color naming stimulus grid used in the World Color Survey (WCS). The grid contains 330 stimulus chips: 320 color chips, and 10 achromatic chips shown in the leftmost column. Participants were shown each chip in a fixed pseudo-random order, and asked to name the color of the chip.

**Figure 2.**
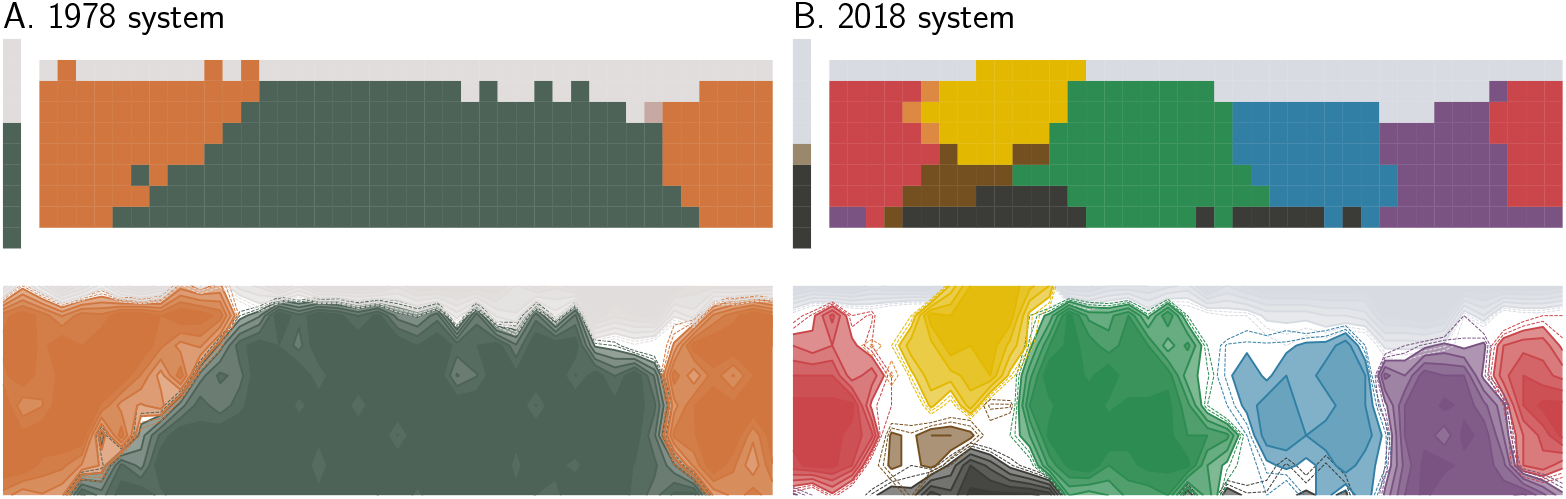
Color naming and its evolution in Nafaanra. The Nafaanra color naming system in 1978 (A) and in 2018 (B), plotted against the color naming grid of Figure 1. Each color term is shown in the color that corresponds to the center of mass of its color category. Mode maps (top) show the modal term for each color chip. Contour plots (bottom) show the proportion of color term use across participants. Dashed lines correspond to agreement levels of 40% *−* 45%, and solid lines correspond to agreement levels above 50%. **(A)** The 1978 system: ‘fiηge’ (light), ‘wͻͻ’ (dark), and ‘nyiε’ (warm or red-like). **(B)** The 2018 system: the three terms from 1978 have smaller extensions and new terms have emerged — ‘wrεnyiηge’ (green), ‘lomru’ (orange), ‘ηgonyina’ (yellow-orange), ‘mbruku’ (blue), ‘poto’ (purple), ‘wrεwaa’ (brown), and ‘tͻͻnrͻ’ (gray).

Our initial data collection began with a pilot study in 2017. Data were collected for Nafaanra by one of us (K.G.), in the same town, Banda Ahenkro, and strictly following the same protocol, which discourages using terms that specify the source of the color, e.g., terms that could be translated into phrases like *fresh leaf*. In the context of Nafaanra in 2017, this effectively meant that participants were restricted to using the original three color terms from the 1978 study: ‘fiηge’, ‘wͻͻ’, and ‘nyiε’. To our surprise, and in contrast to the 1978 data, we found that participants were unable to name a large proportion of the chips when restricted to these three terms, and they expressed frustration at being asked to do so. This suggests a qualitative change in Nafaanra color naming over the recent past. For this reason, subsequent data collection used a free response method, in which no constraints were placed on the color terms that could be supplied as responses. In 2018, 40 years after the original WCS data collection, Nafaanra color naming data were collected again by one of us (K.G.), in the same town, Banda Ahenkro, and following the same protocol, with the exception that participants responded freely in naming the color chips.^2^ Speakers were asked to provide a color term for each chip in the stimulus grid (‘ηga wͻͻ yi hin?’; What is the color?). A total of 15 Nafaanra speakers participated in the 2018 study, 6 female and 9 male, ranging in age from 18-77.^3^

Based on these data, we estimated the 2018 Nafaanra color naming system (Figure 2B) by averaging the naming responses across participants (see Appendix A for individual color naming maps and age data). The 2018 system contains the same three color terms as the 1978 system: light (‘fiηge’), dark (‘wͻͻ’), and warm or red-like (‘nyiε’)—but these now have smaller extensions, and the system also includes seven new color terms: green (‘wrεnyiηge’), orange (‘lomru’), yellow-orange (‘ηgonyina’), blue (‘mbruku’), purple (‘poto’), brown (‘wrεwaa’), and gray (‘tͻͻnrͻ’). While these terms represent the most common responses, there was also some variability in term usage for a few categories; specifically, a small number of speakers used ‘nyanyiηge’^4^ instead of ‘wrεnyiηge’ for green, ‘ndemimi’ or ‘mimi’ instead of ‘ηgonyina’ for yellow-orange, and ‘tra’ instead of ‘wrεwaa’ for brown. One additional term, ‘grazaan’ for red-brown, was used by a single speaker and for a small portion of chips. A more detailed discussion of the terms themselves and how they relate to Twi and English is included in the discussion section.

As can be seen in Figure 2, the Nafaanra color naming system changed substantially between 1978 and 2018, becoming more semantically fine-grained through the addition of new color terms and adjustment in extension of previously existing terms. However, these qualitative observations alone do not determine whether the system has changed in a way that is consistent with pressure for efficiency. To address that question, we turn next to a formal theoretical framework that captures the idea of communicative efficiency and generates precise testable predictions for how color naming may change continuously over time.

## 3. Theoretical framework and predictions

It has been argued that systems of semantic categories are shaped by functional pressure for communicative efficiency (see Kemp et al., 2018, for a review). This general proposal has been explored in the case of color naming (Lindsey et al., 2015; Regier et al., 2015; Gibson et al., 2017; Zaslavsky et al., 2018; Conway et al., 2020), as well as in other semantic domains, such as kinship (Kemp and Regier, 2012), numeral systems (Xu et al., 2020), and indefinite pronouns (Denic et al., 2020). We are interested in testing whether color naming, and semantic systems more generally, change over time while maintaining communicative efficiency.

To this end, we consider the theoretical framework of Zaslavsky et al. (2018, ZKRT), who argued that languages achieve communicative efficiency by compressing meanings into words via the Information Bottleneck (IB) optimization principle (Tishby et al., 1999). This framework is particularly useful in our context for several reasons. First, it is comprehensively grounded in rate-distortion theory (Shannon, 1959; Berger, 1971), the subfield of information theory characterizing efficient data compression under limited resources, offering firm and independently motivated mathematical foundations. Second, it has previously been applied to color naming and was shown to account for much of the known variation across languages, including fine-grained details such as soft category boundaries and patterns of inconsistent naming (Zaslavsky et al., 2018). At the same time, this framework is not specific to color and has also been applied to other semantic domains (e.g., Zaslavsky et al., 2019c), suggesting it may characterize the lexicon more broadly.

Third, this framework provides quantitative predictions not only for the efficiency of attested semantic systems, but also for how they may evolve over time and extend beyond those stages already observed. Specifically, this framework suggests an idealized continuous trajectory of semantic evolution in which efficient systems evolve through gradual adjustments of a single complexity–accuracy tradeoff parameter. In the context of color naming, this theoretically-derived evolutionary trajectory was shown by ZKRT to synthesize key aspects of seemingly opposed accounts of color naming evolution (Berlin and Kay, 1969; MacLaury, 1997; Lyons, 1995; Levinson, 2000). This finding suggests that the ZKRT account may explain substantial aspects of language change. However, that possibility has not yet been tested against diachronic data.

Next, we review ZKRT’s theoretical framework and its predictions, focusing specifically on its instantiation for color naming which we refer to as the IB color naming model.^5^ In Section 4, we will test the predictions of this model on the diachronic Nafaanra color naming data described in the previous section, and assess whether efficiency can explain semantic change over time in Nafaanra.

### 3.1. Communication model

The theoretical framework we review here is based on a simple communication setting (Figure 3A), that can be derived from Shannon’s communication model (Shannon, 1948). Here, we focus on the case in which a speaker and a listener communicate about colors, and attention is restricted specifically to the colors shown in Figure 3B, each of which is represented as a point *U* in a standard perceptual color space, CIELAB. The speaker has a mental representation *M* of one of these colors *U*, drawn from a prior distribution *p*(*m*).^6^ This mental representation *M* is assumed to be a Gaussian distribution in CIELAB space, centered at *U*, capturing the speaker’s mental uncertainty about the color. The speaker communicates this representation by encoding it into a word *W* according to a conditional distribution *q*(*w*|*m*), which serves as a stochastic encoder. The listener receives *W* and attempts to infer from it the speaker’s representation *M* by constructing another representation, 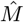, that approximates *M*. The listener’s inferences are Bayesian with respect to the speaker.^7^

**Figure 3.**
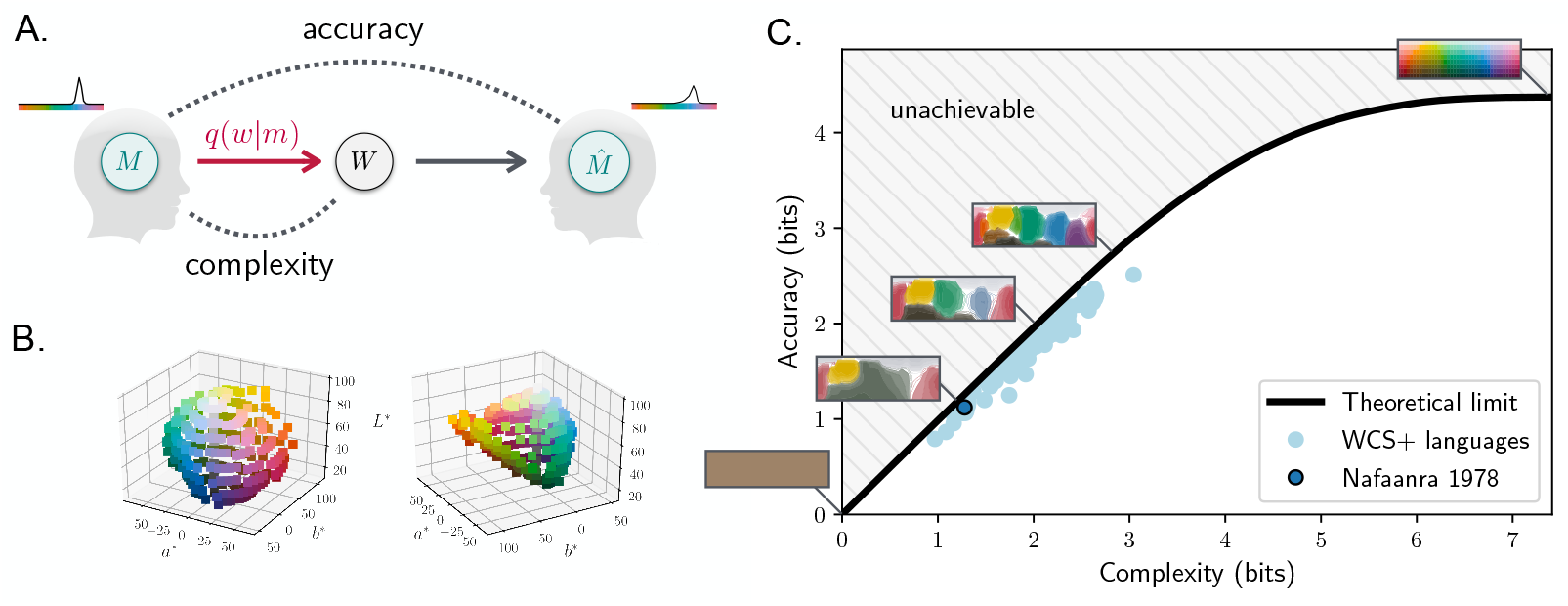
The IB color naming model (adapted from Zaslavsky et al., 2018). **A**. The communication model described in Section 3.1. **B**. Two views of the color stimuli of Figure 1 represented in the three perceptual dimensions of the CIELAB space: *L*^*^ for lightness; (*a*^*^, *b*^*^) for hue and saturation in polar coordinates. **C**. The theoretical limit of efficiency for color naming (black curve) is defined by the set of optimal IB systems for different complexity-accuracy tradeoffs. Contour plots show a few examples of these optimal systems along the curve. Tradeoffs above the curve are unachievable. WCS+ languages include the WCS languages and English. Color naming across languages, including in Nafaanra of 1978, is near-optimal.

That is, given a word *w*, the listener’s inference is defined by

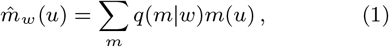

where *q*(*m*|*w*) is obtained by applying Bayes’ rule with respect to *q*(*w*|*m*) and *p*(*m*).

### 3.2. The theoretical limit of semantic efficiency

In this formulation, human semantic systems, such as the Nafaanra color naming systems shown in Figure 2, correspond to encoders *q*(*w*|*m*). The IB principle characterizes the set of optimal systems in this setting, which are parametrized by a single parameter that controls the tradeoff between the complexity and accuracy of the system. As in rate-distortion theory, complexity is measured by the mutual information between the speaker’s mental representation *M* and word *W*,

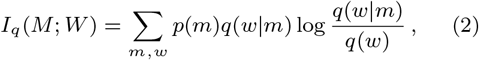

which tightly approximates the number of bits required for communication (Shannon, 1959; Berger, 1971). Accuracy corresponds to the similarity between the speaker’s and listener’s representations, and is measured by *I*_*q*_(*W* ; *U*). Maximizing this second informational term amounts to minimizing the expected Kullback–Leibler (KL) divergence between *M* and 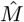 (Tishby et al., 1999; Gilad-Bachrach et al., 2003, and see Appendix B for a detailed derivation in our context). Thus, high accuracy implies that the listener’s inferred representation is similar to the speaker’s representation.

Achieving high accuracy requires a complex lexicon, while reducing complexity may result in accuracy loss. According to the IB principle, optimal systems minimize complexity while maximizing accuracy for some tradeoff *β* ≥ 0 between these two competing objectives. Formally, an optimal encoder *q*(*w*|*m*) for a given value of *β* is one that attains the minimum of the IB objective function,

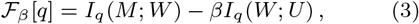

across all possible encoders. Let 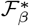 be the minimal value of this objective for a given value of *β*. The theoretical limit of efficiency, also known as the IB curve, is then determined by the set of encoders *q*_*β*_(*w*|*m*) that attain 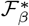 for different values of *β*. This limit in the case of color communication is shown by the black curve in Figure 3C, accompanied by a few examples of optimal encoders along the curve.

### 3.3. Evolution of the optimal systems

Intuitively, the tradeoff parameter *β* controls the relative importance of maximizing accuracy over minimizing complexity, and thus how fine-grained a semantic system is. For *β* ≤ 1, complexity is more important than accuracy, yielding at the optimum a minimally complex yet non-informative system that can be implemented with a single word. This system lies at the origin of the IB curve, as can be seen in Figure 3C. As *β* gradually increases from 1 to ∞, the optimal systems evolve in an annealing process along the IB curve, becoming more complex and more accurate. In general, the optimal systems can also change via reverse-annealing, i.e., when *β* gradually decreases, in which case they will travel down the curve and become less complex. Along this continuous trajectory, the optimal systems undergo a sequence of structural phase transitions at critical values of *β*, in which the number of categories effectively changes (Zaslavsky, 2020).

In the domain of color naming, this theoretical evolutionary trajectory was previously derived from the IB color naming model shown in Figure 3. By mapping the color naming systems of 111 languages (WCS+ dataset) — 110 from the WCS and American English from Lindsey and Brown (2014) — onto optimal systems along this trajectory, it was shown that all of these languages are near-optimal in the IB sense, and that much of the observed cross-language variation can be explained by varying *β* alone. Furthermore, it was shown that the optimal trajectory synthesizes aspects of seemingly opposing accounts of color naming evolution. Berlin and Kay‘s (1969) discrete evolutionary sequence is largely captured by the structural phase transitions that occur at critical points along the trajectory. However, this trajectory is continuous, categories change gradually with *β*, and new ones typically emerge in regions of color space that are inconsistently named. These phenomena resonate with other approaches to the evolution of color naming (MacLaury, 1997; Lyons, 1995; Levinson, 2000) that traditionally appeared to challenge Berlin and Kay‘s (1969) proposal.

As noted by ZKRT, these findings suggest that semantic systems, and color naming in particular, evolve under pressure to remain near the IB theoretical limit and that the optimal evolutionary trajectory, while idealized, may capture substantial aspects of language change. From this perspective, the relative importance of accuracy versus complexity, captured by *β*, may change over time, driving a system up or down along the theoretical limit, but leaving it near-optimal. Thus, this model makes testable predictions for language change.

### 3.4 Quantitative predictions

We adopt the quantitative predictions and evaluation methods derived by ZKRT, and extend them by explicitly considering the dimension of time. If human semantic systems evolve under pressure to be efficient, i.e., to reach the optimum of (3), then the following two properties should hold over time.

*Near-optimality*. For each language *l* with system 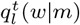 at time *t*, there should be a tradeoff *β*_*l*_(*t*) for which the system is near-optimal. Formally, this means that its deviation from optimality,

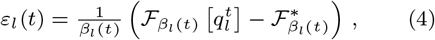

should be small. Because we do not know the true tradeoff parameter, we consider the candidate that maps each system to the nearest point along the theoretical limit, i.e., we take 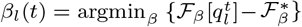. The system 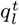 is then taken to be efficient to the extent that *ε*_*l*_(*t*) is small, and this can be assessed with respect to counterfactual data, as described in Section 4. We do not expect *ε*_*l*_(*t*) = 0 because the model does not incorporate every possible factor that may shape language and its evolution. Therefore, we expect that actual systems would only be near-optimal, in the precise sense defined above. For the same reason, transient deviations from optimality are also possible in theory. Our prediction is that large deviations from optimality would not be stable states that are likely to be observed if languages are indeed attracted to the theoretical limit of efficiency.

#### Structural similarity

Considering *ε*_*l*_(*t*) alone reduces the system to only two features — its complexity and its accuracy. However, IB also generates predictions for the full probabilistic structure of 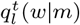. That is, we expect that the full structure of 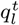 will be similar to that of an optimal system. For simplicity, we compare 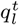 with 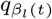, the optimal system at *β*_*l*_(*t*), but note that it is in principle possible that optimal systems at other values of *β* could be more structurally similar to 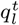. To measure the structural similarity between two probabilistic category systems, we use the generalized Normalized Information Distance (gNID: Zaslavsky et al., 2018) which was designed for this purpose. That is, 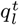 and 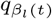 are similar to each other to the extent that the gNID between them is small. In this case as well, we will assess the degree of similarity (1−gNID) relative to counterfactual data.

## 4. Efficiency and language change

The previous work reviewed in Section 3 moved from a synchronic efficiency analysis based on cross-language data to a diachronic hypothesis that language change is shaped by pressure for efficiency. That diachronic hypothesis has not yet been directly tested using fine-grained diachronic data, and the Nafaanra data reported above allow us to fill that gap.

### 4.1. Efficiency over time

First, we are interested in testing whether the efficiency of the Nafaanra color naming system has persisted over time. Because the 1978 Nafaanra data were part of the WCS, we already know from ZKRT’s analyses that the 1978 Nafaanra data lay near the IB limit of efficiency. We conducted an entirely analogous analysis on the 2018 Nafaanra data. Figure 4 shows that the complexity and accuracy of both the 1978 and the 2018 Nafaanra systems are near the theoretical bound, but at different places along the curve. Importantly, Figure 4 and Appendix C show that the 2018 Nafaanra system differs from the English color naming system (estimated from the data of Lindsey and Brown, 2014) and is more efficient than systems obtained by a mixture of the English and 1978 systems. This suggests that pressure for efficiency has shaped Nafaanra beyond its evident contact with English. Thus, these diachronic data from Nafaanra appear to be consistent with the near-optimality prediction.

**Figure 4.**
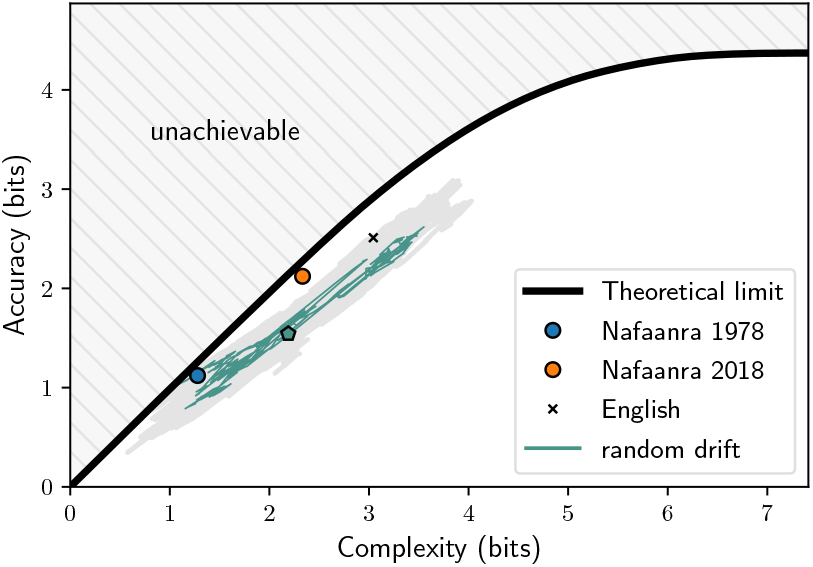
Diachronic efficiency analysis. Color naming in Nafaanra has changed from 1978 to 2018 by climbing up the IB theoretical limit (black curve, same as in Figure 3C). Despite exposure to English, the 2018 Nafaanra system appears at a different tradeoff from the English color naming system, reflecting a qualitative difference between the two systems. The gray area below the curve shows the area covered by 50 hypothetical trajectories traced out by a process of random drift, which were all initialized near the 1978 system. The green trajectory corresponds to the example of Figure 11 in Appendix E, and the pentagon marks its location after 1,500 iterations.

Figures 5A-D compare these two natural systems with their corresponding optimal systems that lie directly on the IB curve. It can be seen that the optimal systems capture substantial aspects of the empirical data, but also differ from those data in some respects. For example, the 1978 system lacks a yellow category that is found in the corresponding optimal system, and the 2018 system has purple and brown categories, while the corresponding optimal system does not. While the early yellow category seems to represent a discrepancy of the model (Zaslavsky et al., 2018), the absence of purple and brown does not necessarily. These categories emerge at a slightly higher value of *β* (see for example Figure 3C), and therefore this mismatch between the model and data may stem simply from noise in our estimation of *β*.

**Figure 5.**
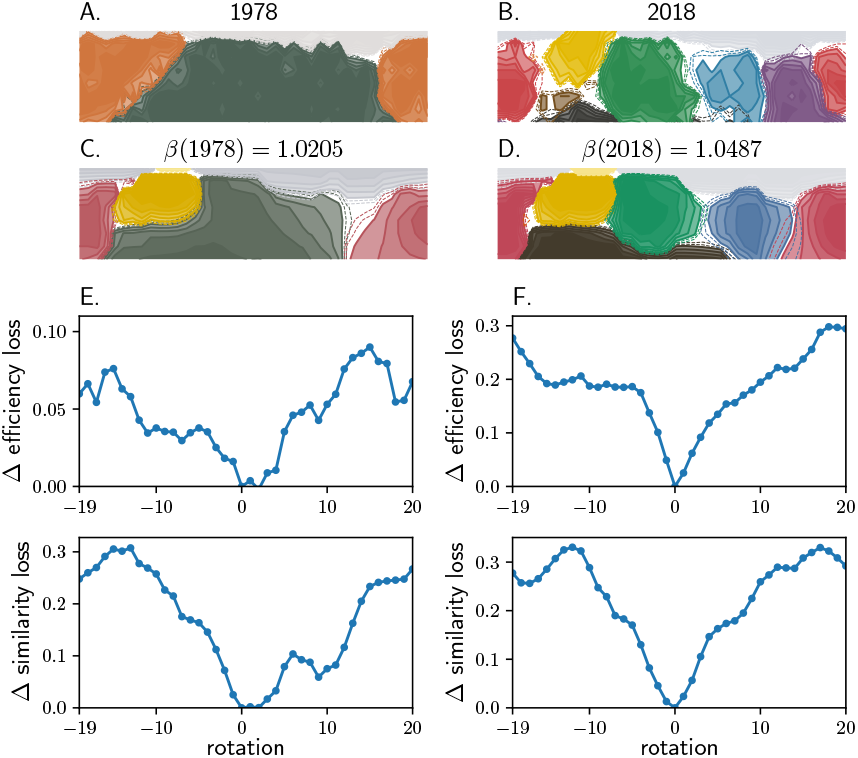
**(A-B)** Empirical data for Nafaanra in 1978 and 2018 (same as Figure 2). **(C-D)** Optimal IB systems corresponding to the actual 1978 and 2018 systems. **(E-F)** Rotation analysis for the 1978 and 2018 Nafaanra systems respectively. Δ efficiency/similarity loss corresponds to the difference between the score of the rotated and actual system (positive values correspond to higher losses of the rotated system).

To quantitatively test the extent to which our predictions hold, we evaluated the efficiency loss (*ε*_*l*_) and similarity loss (gNID) of the 1978 and 2018 systems, and assessed each system with respect to a set of hypothetical variants. These variants were obtained by rotation in the hue dimension (columns of the WCS stimulus grid; Regier et al., 2007) as illustrated in Appendix D, Figure 10. Following ZKRT, in this analysis *β* was fitted to each system separately in order to consider the best scores these hypothetical systems can achieve. Consistent with ZKRT’s findings for the WCS+ languages, including the 1978 Nafaanra system (Figure 5E), the actual (unrotated) 2018 Nafaanra system scores better than any of its hypothetical variants on both measures (Figure 5F). This suggests that the 1978 and 2018 are locally optimal within their set of hypothetical variants, and thus non-trivially efficient. In addition, it can be seen by looking ahead to Figure 6A that these two systems do not deviate much from optimality (less than 0.2 bits), comparable to the average deviation across the WCS+ languages. These results show that 1978 and 2018 Nafaanra are near-optimally efficient when assessed by the same standards that ZKRT used for other color naming systems.

**Figure 6.**
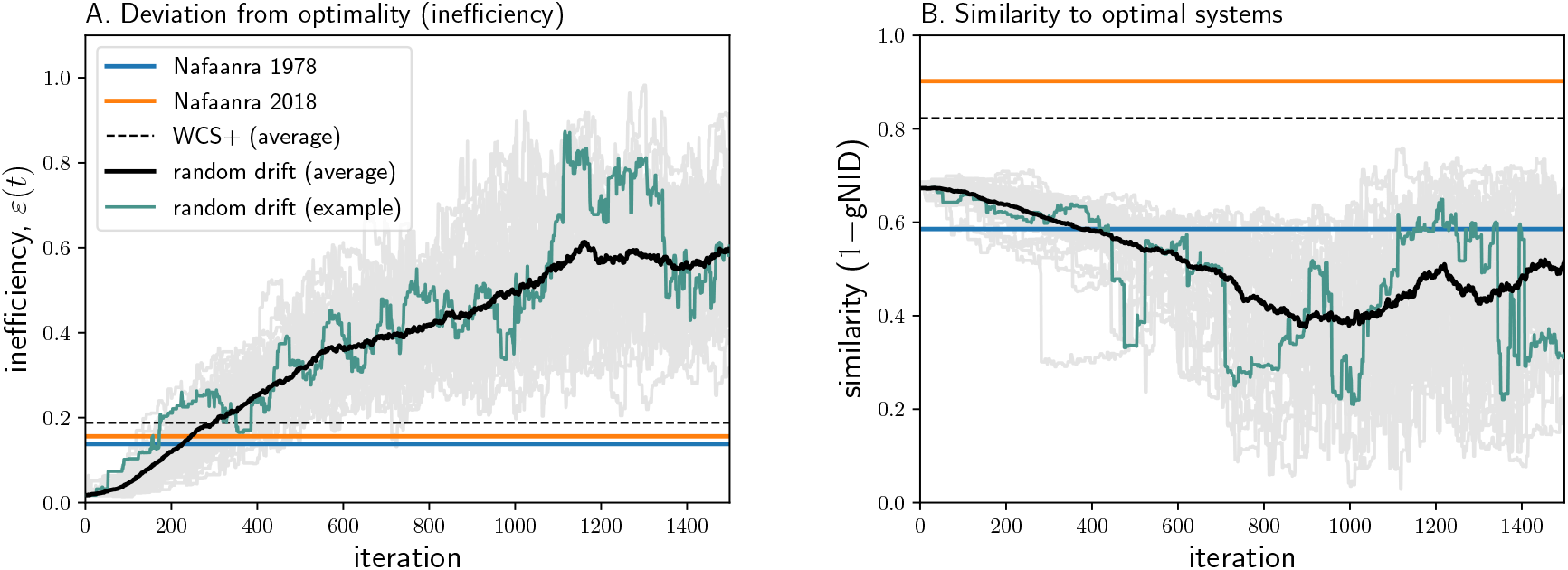
Random drift. **(A)** Deviation from optimality under random drift (lower values are better). Gray curves correspond to 50 hypothetical trajectories. The green trajectory corresponds to the example of Appendix E, Figure 11, and the black curve is the average across all 50 trajectories. Nafaanra’s deviation from optimality is shown by the blue (1978) and orange (2018) horizontal lines. The black dashed horizontal line shows the average deviation from optimality across the WCS+ languages. **(B)** Analogous plot showing the structural similarity (higher values are better) with respect to the corresponding optimal system along the IB curve.

### 4.2. Random drift

So far, we have seen that over the past several decades, the Nafaanra color naming system has changed substantially, while remaining near the theoretical limit of efficiency. This outcome is consistent with our hypothesis that language change may be shaped by functional pressure for efficiency. But before reaching that conclusion, we need to consider a natural alternative: that the same outcome could have been produced by a process of random drift, without any pressure for efficiency. The importance of considering a null model of random drift has recently been emphasized in the literature (e.g. New-berry et al., 2017; Bentz et al., 2018; Karjus et al., 2020), and so here we ask whether a process of random drift could have produced the 2018 Nafaanra system from the 1978 system.

We considered a process of random drift that is described in detail in Appendix E. To avoid random systems, which form a weak baseline, this process maintains some reasonable category structure by representing a color naming system in terms of a set of Gaussian distributions over CIELAB space. It then evolves in a stochastic process that allows existing categories to drift, new categories to emerge, and old categories to occasionally vanish. We generated a set of 50 random drift trajectories, in each case simulating this process for 1, 500 iterations. The initial system was the same for all trajectories, and was obtained by fitting to 1978 Nafaanra, yielding a good approximation of the 1978 system.

The green trajectory in Figure 4 corresponds to one such random drift trajectory, illustrated in Appendix E, Figure 11. The gray area below the IB curve in Figure 4 shows the area traced out by all 50 hypothetical random drift trajectories. It can be seen that these trajectories tend to diverge away from the IB curve, and none reaches the 2018 Nafaanra system. Figure 6A plots the inefficiency (*ε*_*l*_) of the systems in these random drift trajectories over time, and confirms that they tend to become less efficient with time. Interestingly, the same plot also shows that the starting point for these trajectories — a Gaussian approximation to the 1978 Nafaanra system — is more efficient than the 1978 Nafaanra system itself. This demonstrates that the model at the heart of this random drift process can in principle represent highly efficient systems. At the same time, however, the process does not tend to remain at such systems. Figure 6B analogously plots the structural similarity between each system in these trajectories on the one hand, and the corresponding optimal system on the other. It can be seen that the random drift process tends to lead to systems that are dissimilar from those along the theoretical efficiency limit. Given these inefficiency and dissimilarity results, it seems unlikely that this process of random drift could have produced the 2018 Nafaanra system, starting from the 1978 system.

## 5. Discussion

The starting point for this study was the claim that systems of semantic categories evolve under functional pressure for efficiency. This claim is consistent with a substantial amount of synchronic data, but it had not previously been tested directly, by bringing it into contact with fine-grained diachronic data that documents language change over time. The present study has addressed that open issue, by considering the evolution of color naming in Nafaanra over the past several decades, through the lens of efficiency.

We have seen that color naming in Nafaanra has changed substantially while remaining near-optimally efficient, as predicted by the Information Bottleneck (IB) optimality principle and the theory of compression more generally. We have also seen that this outcome would be unlikely under a process of random drift that maintains structured categories but does not incorporate pressure for efficiency. Thus, in at least one language, in at least one semantic domain, and over at least one stretch of time, it appears that a semantic system has evolved in a way that reflects functional pressure for efficiency. However, the information-theoretic framework we have employed in this work and its predictions for language change are not specific to these settings. In fact, this framework has recently gained substantial cross-linguistic support in several other domains, including container naming, animal taxonomies, personal pronouns, and grammatical number, tense and evidentiality (Zaslavsky et al., 2019c, 2021; Mollica et al., 2021). However, as in the case of color naming, these results have so far been based mainly on synchronic data. Therefore, an important direction for future research is to further test the diachronic predictions of this theory in more languages, domains, and periods of time. Interestingly, this framework can also be used to study the influence of communicative need on language change. In this evolutionary view of language, communicative need parameterizes the IB objective function (Zaslavsky et al., 2019a), which in turn, serves as a fitness criterion guiding the ways in which systems of semantic categories change.

Our findings converge with those of a complementary line of work. In a comment on the finding that systems of semantic categories tend to be efficient, Levinson (2012) asked “where our categories come from” – i.e. what process gives rise to these efficient category systems. He suggested that some insight into this question might be obtained from studies of iterated learning that simulate language evolution in the lab (e.g. Kirby et al., 2008; Xu et al., 2013). This suggestion inspired Carstensen et al. (2015) to explore whether simulated language evolution in the lab in fact produces systems of increasing efficiency. They found that it does, and more recent work has probed these ideas more closely (Carr et al., 2020). Although these earlier studies were based on different formulations of the notion of efficiency, the present work resonates with their findings by showing that actual language change, not just simulated language change, tends toward communicatively efficient semantic systems. More recently, Chaabouni et al. (2021) showed that artificial neural agents playing a cooperative color-discrimination game develop color signaling systems that converge to the same IB theoretical limit of efficiency that was proposed by ZKRT and considered in this work. This suggests that the computational principles underlying language change in humans may be crucial for evolving human-like communication in artificial agents.

At the same time, the present findings leave a number of points open, some of which suggest additional directions for future research. We have considered a specific model of random drift for category systems, and while we believe this model to be a reasonable one, it is conceivable that other models of drift could yield different results. More fundamentally, although we have spoken of language evolving under pressure for efficiency, and although our findings are consistent with that idea, we do not know the shape of the trajectory that took Nafaanra from where it was in 1978 to where it was in 2018. The evolution we have seen could have come about in a series of small incremental changes, tracing the IB curve closely, or the system could have been pulled fairly far away from efficiency by some external force, such as language contact, and then gradually retreated to efficiency.

Language contact is an especially relevant consideration in the case of Nafaanra, given the exposure of Nafaanra speakers to English and Twi, as noted above (see Huisman et al. (2021) for a comparable situation). While it is not known to what extent the evolution in Nafaanra color naming is attributable to contact, it is possible that some of the new 2018 Nafaanra categories may have been borrowed or calqued. For example the word ‘mbruku’ (blue) may plausibly be a borrowing from English ‘blue’ or from Twi in which ‘bruu’ is sometimes used for blue, though ‘bibire’ is also used. Likewise, the Nafaanra term ‘poto’ (purple) may also be borrowed from English, whereas the Twi word, ‘brεdum’ has minimal phonetic similarity and is likely not a borrowing source. Some other Nafaanra terms, if influenced by another language, seem more likely to have been influenced by Twi than by English. For example ‘ηgonyina’ (yellow-orange) is Nafaanra for chicken fat and ‘wrεnyiηge’ (green) means fresh leaf, reasonable descriptions of the colors involved. In Twi, the terms ‘akokͻsradeε’ (yellow) and ‘ahabammono’ (green) likewise mean chicken fat and fresh leaf, respectively; thus the form of these color terms may be calques from Twi, as there is no phonetic similarity between the terms — or these terms may have developed independently because these referents are locally culturally salient examples of these colors.

Importantly, however, although the 2018 Nafaanra system shares some features with the English and Twi systems, it is not a simple copy of either: the category pink is missing from Nafaanra although it is present in both English and Twi (‘memen’), the category orange is minimal and only barely visible in the contour plot of Figure 2B, and the 2018 system has retained the three named categories of the 1978 Nafaanra system, with the same names but with adjusted extensions. Thus, even if substantial parts of the 2018 Nafaanra system were either borrowed from or motivated by English and/or Twi, some “naturalization” process appears to have occurred whereby the categories adjusted to form a coherent system in Nafaanra — and we have seen that the resulting system is an efficient one. Further work will be needed to more fully ascertain the role of language contact, and, to the extent possible, the details of the historical trajectory of Nafaanra language change relative to the theoretical limit. However, whatever the details of that trajectory, our current results based on the beginning and end points of that trajectory do suggest a process that is in some way constrained to either remain, or eventually return to, near the theoretical limit of efficiency.

The collection of the new Nafaanra color naming data grew out of an informal exchange between two of the authors, K.G. and T.R., in a classroom setting. T.R. was presenting color naming data from the World Color Survey, and K.G., who was taking the class, mentioned that she was very familiar with one of the WCS languages, Nafaanra, because it was a focus of her ongoing linguistic fieldwork. This led naturally to the idea of K.G. collecting new Nafaanra color naming data the next time she returned to the field. With this idea in hand, it actually came as a bit of a surprise to us to realize that the WCS data were now old enough to be of some historical interest. Although the data were collected in the 1970s, they were only digitized and web-posted in the early 2000s, and they continue to be a widely and regularly used data resource — that is, the data “got old” gradually and without anyone remarking on that fact — until the realization we have just mentioned. That realization, and the follow-up work on Nafaanra reported here, open the possibility of analogous follow-up studies for any or all of the 109 other languages in the WCS, to more comprehensively test the hypothesis we have explored: that color naming evolves under pressure for efficiency.

## Acknowledgments

This paper is dedicated to the late Professor Naftali Tishby. Tali was the PhD advisor of Noga Zaslavsky, and the computational analysis in this paper was developed as part of her PhD thesis and was deeply inspired by Tali’s principled scientific approach. Tali was a rare scientist whose research and vision made a profound impact on the understanding of the computational principles that govern both natural and artificial intelligence. He is greatly missed.

We thank Paul Kay for helpful discussions, the Nafaanra community for their help in collecting the data, and Phoebe Killick and Hsin-Yeh Tsai for their help in digitizing the raw data. We also thank Delwin Lindsey and Angela Brown for kindly sharing their English color naming data with us. This study was partially supported by an MIT Brain and Cognitive Sciences Fellowship in Computation (N.Z.), Robert L. Oswalt Graduate Student Support Endowment for Endangered Language Documentation (K.G.), ARC grant FT190100200 (C.K.), and DTRA grant HDTRA11710042 (T.R.).

## Appendix A. Nafaanra 2018 individual maps

To provide a complete view of the 2018 Nafaanra color naming data, we present here the color naming map for each participant in the data (Figure 7). It can be seen that even the participants who were born before 1978 (ages 48–77) exhibit naming patterns that are similar to the 2018 system (Figure 2B) and more refined than the 1978 system (Figure 2A). This supports our claim that color naming in Nafaanra has changed since 1978.

**Figure 7.**
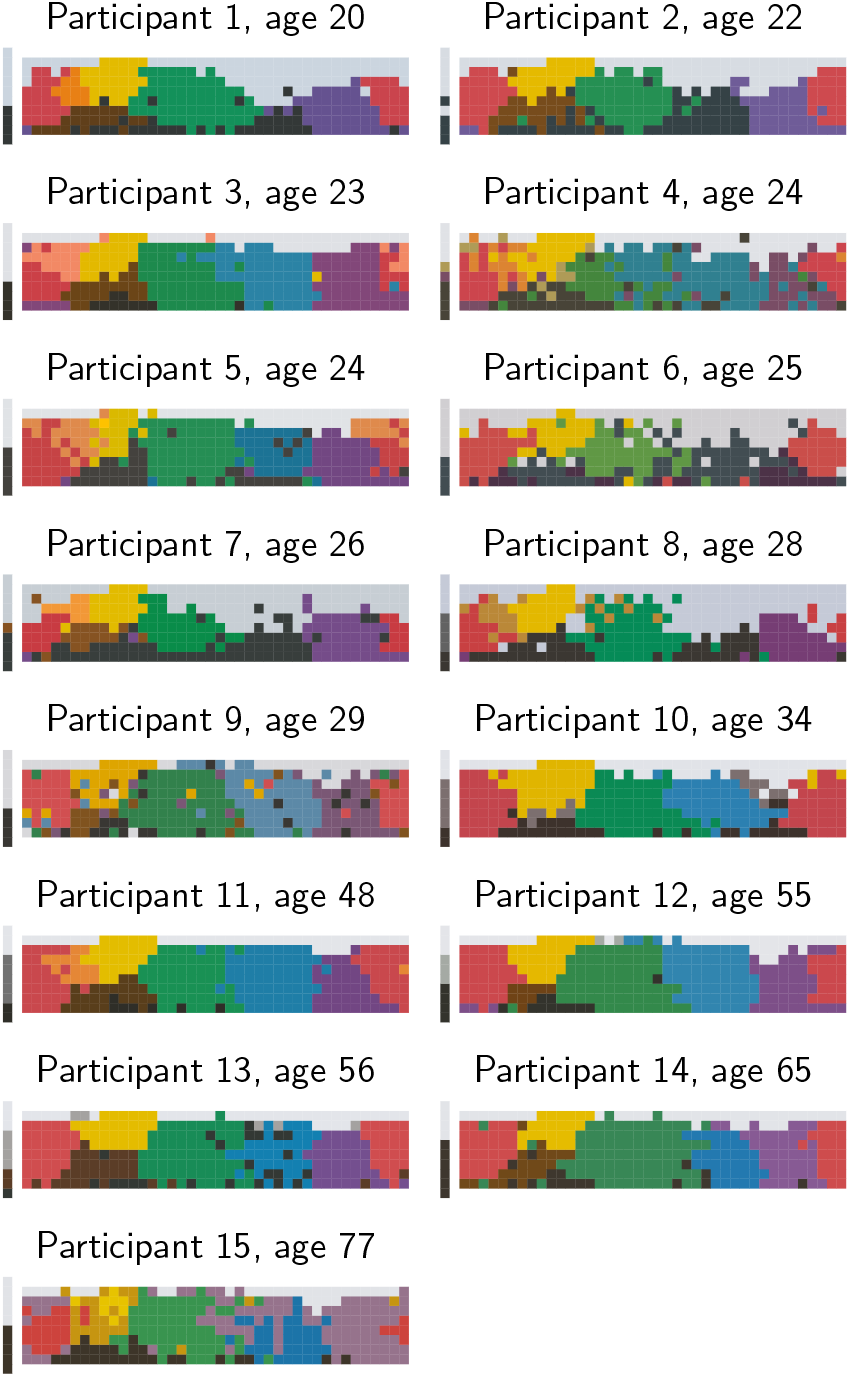
Color naming maps for each participant in the 2018 Nafaanra color naming data. Participants are sorted by age. Each chip in the stimulus grid (Figure 1) is colored according to its term. The color associated with each term is the color centroid of the term’s color category, evaluated per participant.

## Appendix B. Accuracy and distortion in the Information Bottleneck

Section 3 refers to the fact that *I*(*W* ; *U*) corresponds to the similarity between the speaker’s and listener’s representations, and is therefore taken to be the accuracy term in the Information Bottleneck (IB) framework. This has previously been shown for IB in general (Tishby et al., 1999; Gilad-Bachrach et al., 2003), and see (Zaslavsky, 2020) for a detailed discussion of this derivation for the special instantiation of IB for semantic systems. For completeness, we review below the derivation of *I*(*W* ; *U*) as the natural accuracy measure in our setting.

Recall that each speaker meaning is defined by a distribution, or belief, over the domain *U*. Thus, we denote by *m*(*u*) the probability of *u* (in our case, *u* is a color) given that *m* is the speaker’s mental representation. Similarly, 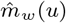 denotes the probability that the listener assigns to *u*, given that the listener infers 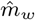 as the mental representation in response to the speaker’s word *w*. The KL-divergence between the speaker’s and listener’s mental representations is defined as

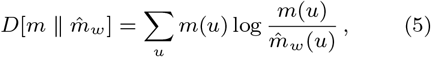

and the total expected divergence (or distortion) is defined as

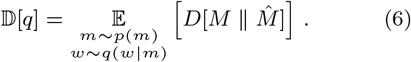

Now, let *m*_0_(*u*) = Σ_*m*_ *p*(*m*)*m*(*u*) be the a-priori mental representation. If the listener’s inferences obey equation (1), as is the case in IB, then the following holds:

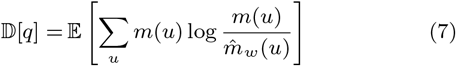

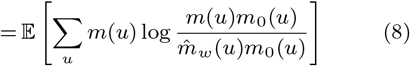

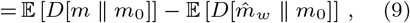

where (9) follows from substituting equation (1) in the expectation of the second term. Note that the first term is constant, namely it does not depend on the speaker’s encoder nor on the listener’s inferences. The second term is an equivalent definition of *I*(*W* ; *U*), and it measures the amount of information that the speaker’s words contains about the speaker’s intended colors. Therefore, minimizing the total divergence 𝔻 [*q*] is equivalent to maximizing *I*(*W* ; *U*), and the latter is the natural measure of accuracy.

## Appendix C. Efficiency beyond contact

Our results in the main text suggest that although exposure to English may have inspired some of the changes in Nafaanra, the 2018 Nafaanra system reflects pressure for efficiency beyond the influence of language contact. Here we provide additional analysis in support of this claim. First, Figure 8 shows that the 2018 system differs from the English system (estimated from the data of Lindsey and Brown, 2014) not only in the number of color categories but also in their extension. This suggests that the 2018 system cannot be explained by simply copying English color categories into Nafaanra.

**Figure 8.**
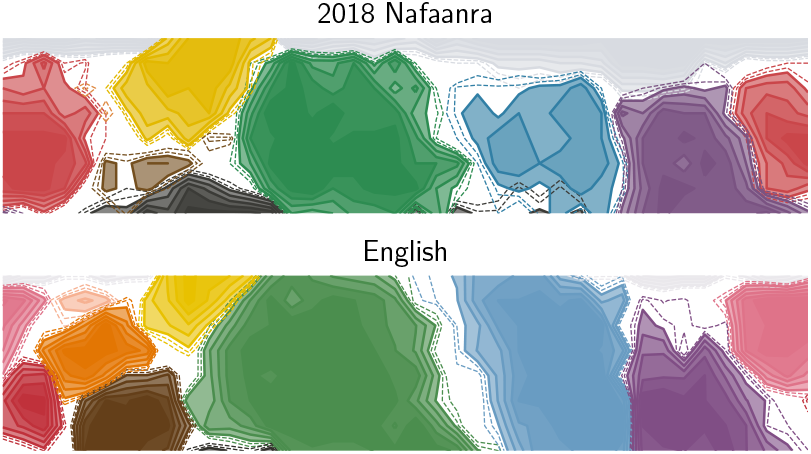
Contour plots of the Nafaanra (2018, same as Figure 2B) and English systems. While the two systems share some resemblance, they differ both in the number of color categories (e.g., pink does not appear in Nafaanra) and in their extensions (e.g., the brown and blue categories are larger in English, while the yellow and black/dark categories are larger in Nafaanra).

Second, we compared our results with a simple baseline model of language contact that does not take into account any pressure for efficiency. Specifically, we considered a set of hypothetical systems that are obtained by linear mixtures of the 1978 system and the English system. Let *P*_78_(*w*|*c*) and *P*_eng_(*w*|*c*) be the empirical distributions of terms *w* given colors *c*, as estimated from the 1978 Nafaanra and English data respectively. In order to combine these systems, we first need to align their categories. To this end, we mapped each term in the 1978 Nafaanra system to its corresponding English term, using the English terms “white,” “black,” “red,” and “gray.”^8^ In addition, to allow the 1978 system to potentially evolve to the full English system, we added hypothetical terms corresponding the remaining English terms but with zero probability mass. In other words, we constructed 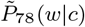 from the 1978 system such that 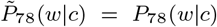 if *w* appeared in 1978 and 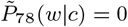 otherwise. We then considered the following set of hypothetical systems:

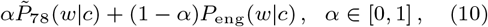

where the 1978 system is obtained at *α* = 1, the English system is obtained at *α* = 0, and in between we get linear mixtures of the two systems.

Figure 9 compares the complexity and accuracy tradeoffs of these hypothetical mixture systems with those of Nafaanra and English and the optimal tradeoffs at the IB theoretical bound. It can be seen that the 2018 system is more efficient (i.e., lies closer to the theoretical bound) than the mixture systems, in addition to being distant from the English system. This further supports our conclusion that Nafaanra has changed under pressure for efficiency beyond the influence of language contact.

**Figure 9.**
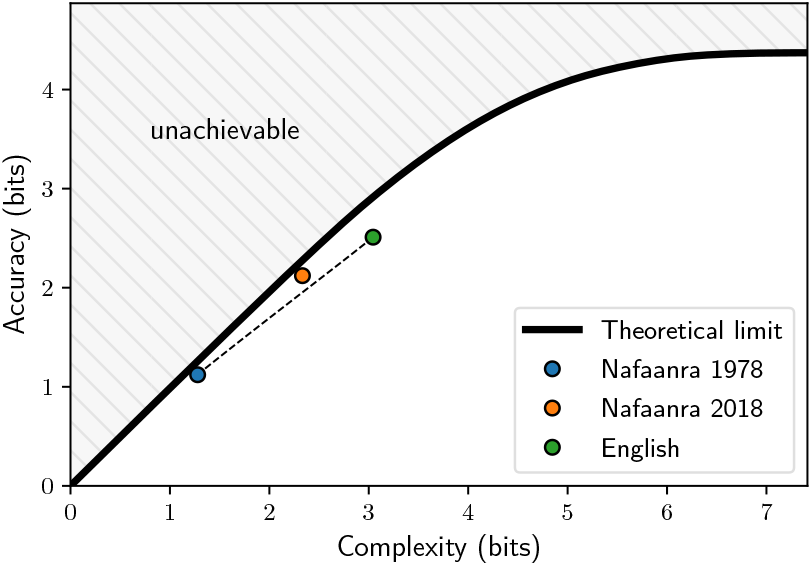
Similar to Figure 4. Dashed line corresponds to the hypothetical systems obtained by mixtures of the 1978 and English systems. The 2018 system is more efficient than these mixtures and does not lie near the English system, suggesting that the changes in Nafaanra were shaped by pressure for efficiency beyond the influence of exposure to English.

## Appendix D. Rotation analysis

Our evaluation of the efficiency of the Nafaanra color naming system with respect to a set of hypothetical systems is based on Regier et al.’s (2007) rotation analysis. That is, for each color naming system, a set of hypothetical systems can be derived by rotations along the hue dimension of the WCS color naming grid (Figure 1). This is illustrated in Figure 10 for the 2018 Nafaanra system.

**Figure 10.**
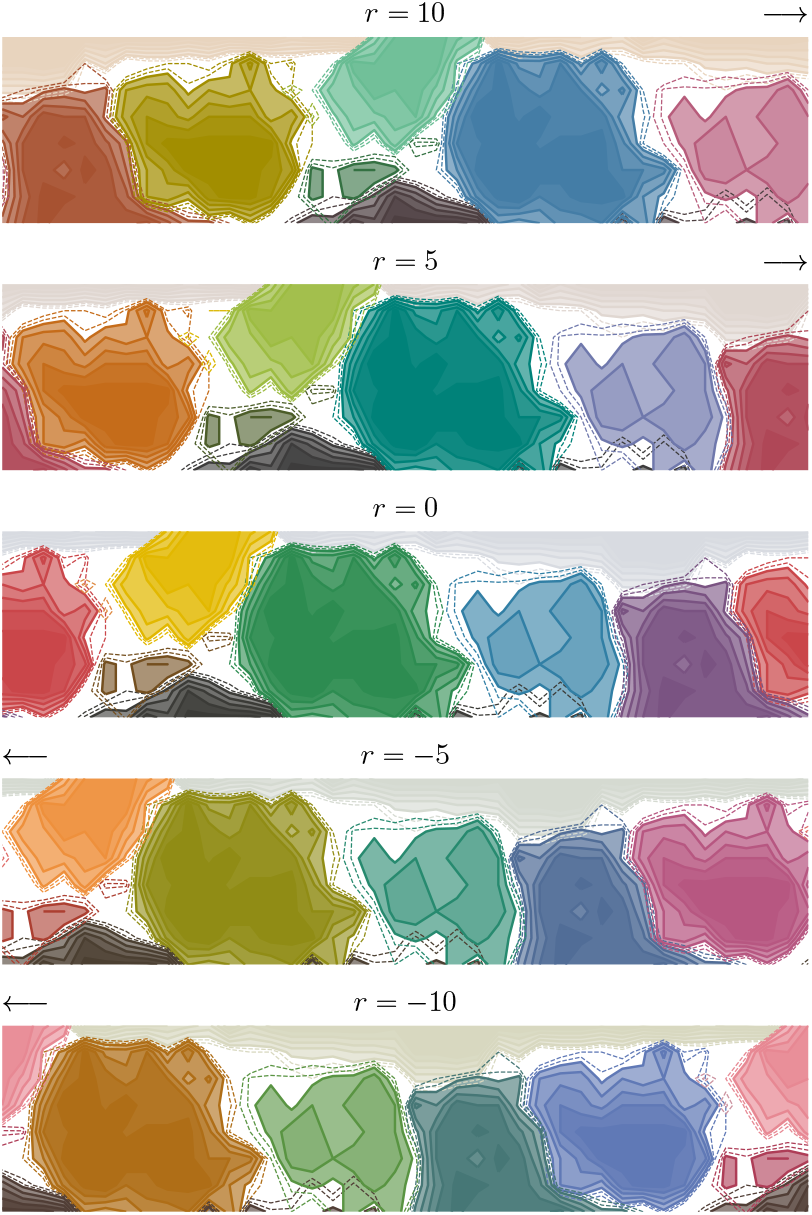
Example of rotated variants of the 2018 Nafaanra system. *r* = 0 corresponds to the actual system, *r >* 0 to a shift of *r* columns to the right with respect to the color grid of Figure 1, and *r <* 0 to a shift of |*r*| columns to the left.

## Appendix E. Random drift model

Our random drift model simulates language change via a stochastic process that preserves structured categories without incorporating pressure for efficiency. To this end, we consider a class of artificial color naming systems, in which each category *w* induces a Gaussian distribution, *q*(*c*|*w*) = η (*c*; *μ*_*w*_, Σ_*w*_), over CIELAB space (Abbott et al., 2016). In practice, we discretized these Gaussians by restricting them to colors of the WCS grid (Figure 1). A system with *k* categories is defined by *k* Gaussians, and a *k*-dimensional probability vector *q*(*w*). Given these parameters, the naming distribution is taken to be *q*(*w*|*c*) ∝ *q*(*c*|*w*)*q*(*w*), where *c* is a color. Our stochastic process takes an initial system from this class, and propagates it in time by allowing its parameters to change gradually.

Before we define the dynamics of this process, our parameterization requires further elaboration. First, to ensure that each covariance matrix Σ_*w*_ remains positive semi-definite, we parameterize it by another matrix, *L*_*w*_, such that 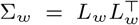. Second, to allow categories to emerge or vanish, we assume a maximum of *K* = 330 potential categories, and keep a weight vector, *π*_*w*_, for them. Only categories for which *π*_*w*_ is higher than a given threshold *η* are included in the lexicon. For those categories, we define *q*(*w*) ∝ *π*(*w*). Therefore, *η* is a hyper-parameter that controls the tendency to add new categories. At the *t*-th iteration of the process, the system is defined by 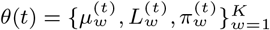.

Given an initial system, *θ*(0), the dynamics of the process are defined as follows. At each iteration *t*, a category *w*_*t*_ is chosen at random. First, the weight vector is updated by randomly selecting whether to add or subtract *η* from 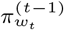, and keeping the vector non-negative and normalized. Next, if *w*_*t*_ is already in the lexicon, i.e. 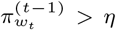, then with probability 0.5 its parameters are updated as follows:

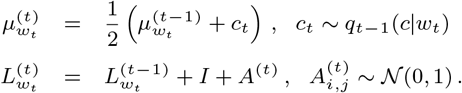

The update rule for 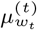 shifts it in the direction of *c*_*t*_, which on average would be a small shift because *c*_*t*_ is sampled from *q*_*t*−1_(*c*|*w*_*t*_). The update rule for 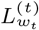 adds to it a noise matrix, *A*^(*t*)^, and the identity matrix, *I*, in order to encourage the category to grow over time.

Finally, it remains to set the initial set of parameters, *θ*(0), and threshold *η*. We set *θ*(0) such that the corresponding system will approximate the actual 1978 Nafaanra system. For each category *w* in the 1978 system, we fit a Gaussian with a diagonal covariance matrix to that category, and take 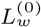 to be its square root. For these categories, we take 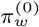 to be their proportion in the 1978 naming data. For the remaining potential categories, which are not in the lexicon: we set 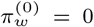, initialize 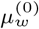 by randomly selecting a chip from the WCS grid (with replacement), and initialize 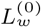 by 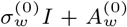, where 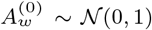 and 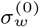 is drawn uniformly from [1, 5]. We take *η* = 0.01, for which we observed a trend of gradual increase in the number of categories, reaching on average *k* = 23.9 after 1, 500 iterations. An example of a hypothetical trajectory that was generated by this random drift process is shown in Figure 11.

**Figure 11.**
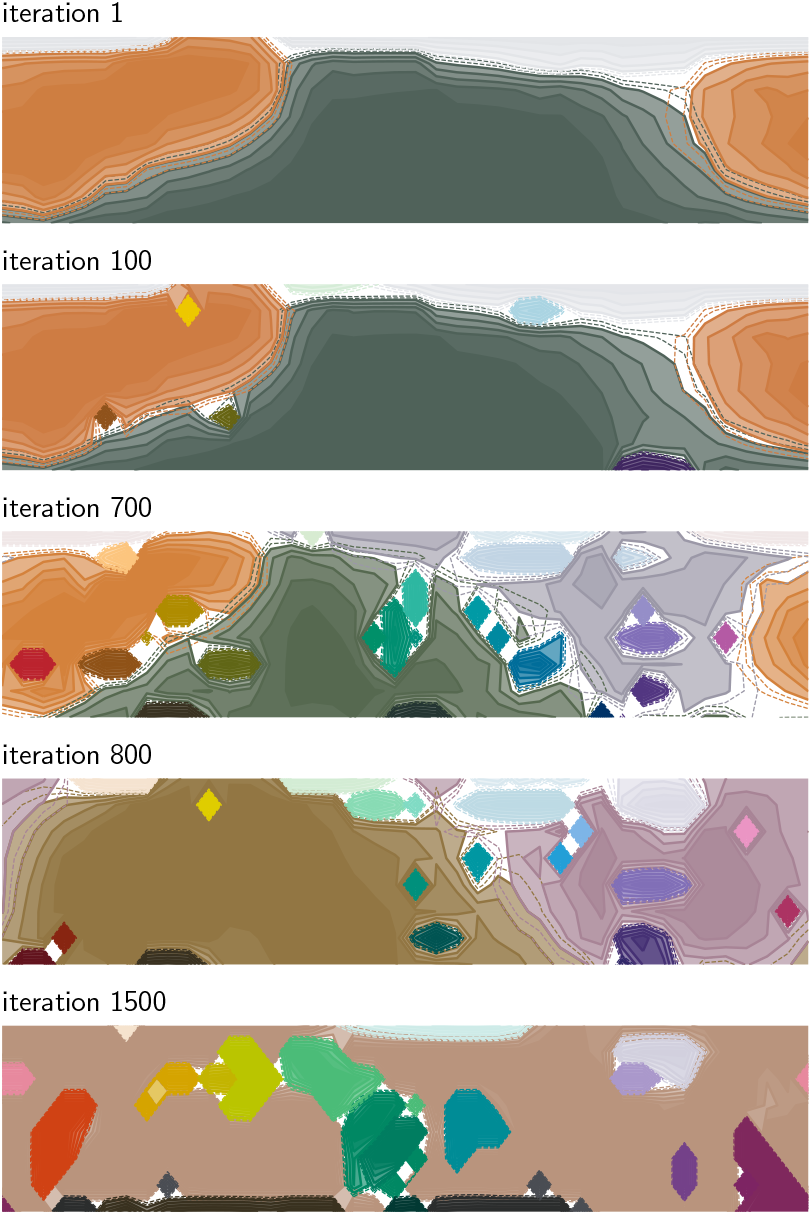
An example of a hypothetical trajectory generated by a process of random random drift. The trajectory was initialized at a system fitted to the 1978 Nafaanra system, and traced out for 1,500 iterations.

## Footnotes

1 WCS data are available at http://www.icsi.berkeley.edu/wcs/data.html. WCS protocol is specified in the Instructions to Fieldworkers, available at https://www1.icsi.berkeley.edu/wcs/images/WCS_instructions-20041018/jpg/border/index.html.

2 For example, following WCS protocol, the 2018 study was conducted on bright days in the shade to ensure chip visibility and data compatibility with the 1978 data. The chips used were the same as those used in 1978, and were presented in the same order.

3 Free-response naming data were also collected in 2017 from 10 participants (6 male and 4 female, ranging in age from 20-68). Our results for the 2017 and 2018 free-response naming data are qualitatively similar.

4 The term ‘nyanyiNge’ only occurs in the 2017 pilot data for a single speaker.

5 The IB color naming model is publicly available at https://github.com/nogazs/ib-color-naming.

6 We take *p*(*m*) to be the prior originally used by ZKRT. See (Zaslavsky et al., 2018, 2019b) for more details about this prior, and (Zaslavsky et al., 2019a) for an evaluation of several alternative priors.

7 This is not an assumption of the model, as it can be derived directly from the IB optimality principle (see Zaslavsky et al., 2018, SI Section 1.2.).

8 We also considered an alignment that is based on map-ping each term in the 1978 system to the English term that has the closest color centroid in CIELAB space. This does not change the results of Figure 9.

